# Conversion of methionine biosynthesis in *E. coli* from trans- to direct-sulfurylation enhances extracellular methionine levels

**DOI:** 10.1101/2023.03.28.534524

**Authors:** Nadya Gruzdev, Yael Hacham, Hadar Haviv, Inbar Stern, Matan Gabay, Itai Bloch, Rachel Amir, Maayan Gal, Itamar Yadid

**Author notes:** **Corresponding authors** Itamar Yadid;, Maayan Gal.

## Abstract

Methionine is an essential amino acid in mammals and a critical metabolite in all organisms. As such, various applications, including food, feed, and pharmaceuticals, necessitate the addition of L-methionine. Although amino acids and other metabolites are commonly produced through bacterial fermentation, high-yield biosynthesis of L-methionine remains a significant challenge due to the strict cellular regulation of the biosynthesis pathway. As a result, methionine is produced primarily synthetically, resulting in a racemic mixture of D,L-methionine. This study aimed to enhance methionine bio-production yields in *E. coli* by replacing its highly regulated trans-sulfurylation pathway with the more common direct-sulfurylation pathway used by other bacteria. To this end, we generated an auxotroph *E. coli* strain (MG1655) by simultaneously deleting *metA* and *metB* genes and complementing them with *metX* and *metY* from different bacteria. Complementation of the genetically modified *E. coli* with *metX/metY* from *Cyclobacterium marinum* or *Deinococcus geothermalis*, together with the deletion of the global repressor *metJ* and overexpression of the transporter YjeH, resulted in a substantial increase of up to 126 and 160-fold methionine relative to the wild-type strain, respectively, and accumulation of up to 700 mg/L using minimal MOPS medium and 2 ml culture. Our findings provide a method to study methionine biosynthesis and a chassis for enhancing L-methionine production by fermentation.

**Highlights:**

- Replacement of *E. coli metA* and *metB* with *metX* and *metY* recovered its growth
- The engineered *E. coli* has a 160-fold increase in extracellular methionine levels
- Selection of different *metX* and *metY* leads to varying growth rates and enhanced methionine levels

## 1. Introduction

Methionine is a sulfur-containing amino acid that is synthesized by plants, fungi and bacteria, but not by vertebrates; thus, it is considered an essential amino acid. Although it is one of the less abundant amino acids in proteins (Pasamontes & Garcia-Vallve, 2006), its hydrophobic nature contributes significantly to the stabilization of proteins’ structure (Bigelow & Squier, 2005; Valley et al., 2012). Methionine also plays an essential role in initiating mRNA translation and indirectly regulates a variety of cellular processes as the precursor of *S*-adenosyl-methionine (SAM), the primary biological methyl group donor (Cantoni, 1953; Gophna et al., 2005). As a necessary amino acid for vertebrates, methionine must be obtained from external sources as a nutrient. However, the low level of methionine in plants limits the nutritional quality of plant-based diets, and external supplementation of L-methionine to food is necessary to achieve the requisite levels for the development of humans and livestock (Carvalho et al., 2018). Microbial production of methionine beyond physiological levels is challenging, due to its high cellular energy demands and the strict cellular regulation of its synthesis and accumulation (Figge, 2006; Becker & Wittmann, 2012; Li et al., 2017). The primary method to produce methionine for food and feed supplementation involves chemical synthesis, resulting in a racemic mixture of D,L-methionine and additional toxic compounds that must be removed from the final product. Thus, there is increasing demand to produce the natural form of L-methionine through an efficient bio-fermentation process (Willke, 2014; Shim et al., 2017; Schipp et al., 2020).

### 1.1 Enhancement of methionine production in *E. coli*

Efforts to produce methionine in bacteria have mainly focused on the clearance of negative regulation, controlling the metabolic flux in and out of the pathway and removing feedback inhibition of enzymes that comprise part of the biosynthesis pathway (Nakamori et al., 1999; Usuda & Kurahashi, 2005; Maier et al., 2009; Li et al., 2017; Sagong et al., 2022). For instance, disruption of *metJ*, which is a master regulator of the methionine pathway, together with overexpression of the genes encoding for MetA and the methionine-exporter YjeH, resulted in an approximately ten-fold improvement in the production of L-methionine in *E. coli* (Liu et al., 2015; Huang et al., 2017). Additionally, enhancing the synthesis of upstream precursors required for methionine biosynthesis, alongside simultaneous alteration of multiple pathways, was shown to be important for an optimized methionine bioproduction (Figge, 2006; Dischert et al., 2015; Huang et al.; 2018). Moreover, modifications of regulatory elements, controlling the expression of multiple related genes, and supplementation of the bacterial growth medium with specific metabolites that were identified as limiting factors all resulted in a substantial increase of methionine levels of up to 18 g/L (McCoy, 2016; Li et al., 2017; Zhou et al., 2019; Tang et al., 2020).

### 1.2 Bacterial direct- and *trans*-sulfurylation pathways for biosynthesis of methionine

Trans- and direct-sulfurylation are the two main pathways for methionine biosynthesis in bacteria. As the names imply, the two pathways differ mainly in the sulfur assimilation steps (Foglino et al., 1995; Vermeij & Kertesz, 1999; Hacham et al., 2003; Ferla & Patrick, 2014; Brewster et al., 2021). In trans-sulfurylation, homoserine is converted to L-homocysteine in three steps that are catalyzed by the enzymes MetA, MetB, and MetC, which are also known as L-homoserine *O*-succinyl transferase (HST; EC 2.3.1.46), cystathionine gamma synthase (CgS; EC 2.5.1.48), and cystathionine beta lyase (CbL; EC 4.4.1.13), respectively (Figure 1). MetA synthesizes *O*-succinyl or *O*-acetyl L-homoserine, and MetB uses cysteine and *O*-succinyl L-homoserine to form cystathionine. In this pathway, inorganic sulfur in the form of hydrogen sulfide is first incorporated into cysteine by the enzyme *O*-acetylserine sulfhydrylase A (CysK), such that cysteine serves as the sulfur donor for the following synthesis of methionine (Kawano et al., 2018). MetC converts cystathionine into L-homocysteine.

**Figure 1.**
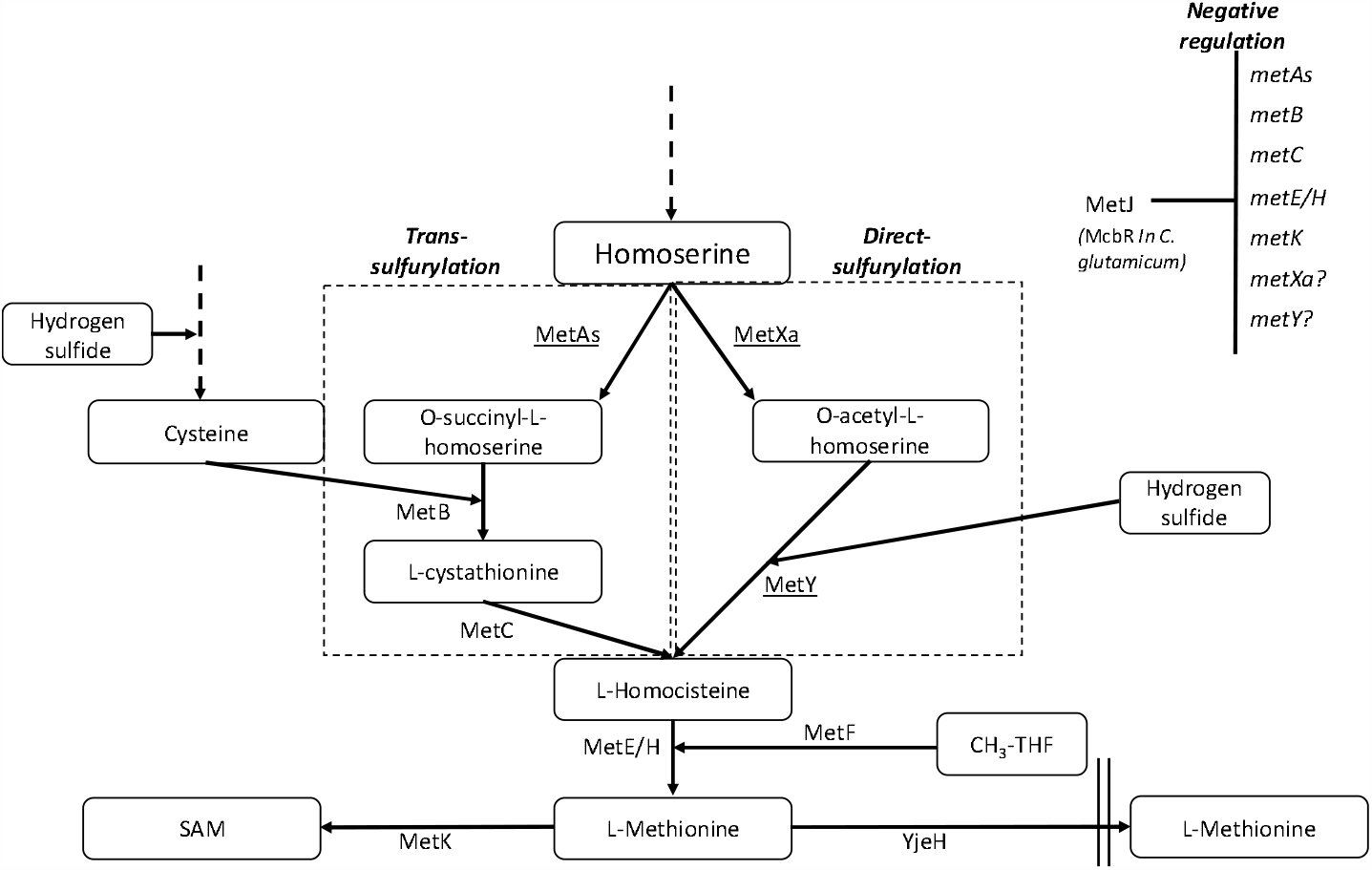
Direct- and trans-sulfurylation of methionine biosynthesis in bacteria. The first step in methionine biosynthesis involves the activation of homoserine through an acylation step. Two enzymes encoded by *metA*s and *metX*a genes (Bastard et al., 2017) activate homoserine. The enzyme homoserine succinyl transferase (HST, MetAs) converts homoserine and succinyl-CoA into *O*-succinyl-L-homoserine (OSH). The enzyme homoserine acetyl transferase (HAT, MetXa) converts homoserine and acetyl-CoA into *O*-acetyl-L-homoserine (OAH). In the trans-sulfurylation pathway, cysteine and *O*-succinyl-L-homoserine (OSH) are converted into cystathionine by cystathionine-γ-synthase (CgS, MetB). Cystathionine is converted into homocysteine by cystathionine-β-lyase CbL (MetC). In the direct-sulfurylation pathway, OAH is converted into homocysteine by the O-acetylhomoserine sulfhydrylase (OAHS, MetY). Metabolites in the pathways are boxed. MetJ and McbR are master negative regulators in *E. coli* and *C. glutamicum*, respectively. Additional abbreviations: MetF -5,10-methylenetetrahydrofolate reductase, MetK - S-adenosylmethionine synthase, MetE - Cobalamin-independent methionine synthase, MetH - Methionine synthase, YjeH - L-methionine exporter.

In the direct-sulfurylation pathway, L-homoserine is converted into L-homocysteine in only two steps, catalyzed by the enzymes MetX and MetY, known as L-homoserine *O*-acetyltransferases (HAT; EC 2.3.1.31) and *O*-acetylhomoserine sulfhydrylase (OAHS; EC 2.5.1.49), respectively. MetX produces *O*-acetyl L-homoserine from homoserine and acetyl-CoA, while MetY combines *O*-acetyl L-homoserine with an inorganic sulfur in the form of hydrogen sulfide to form L-homocysteine; thus, the latter does not rely on cysteine as the sulfur source (Figure 1).

### 1.3 Exploring alternative pathways for sulfur assimilation of methionine in *E. coli*

While most bacteria use the direct-sulfurylation pathway for methionine biosynthesis, *E. coli* utilizes the trans-sulfurylation (Weissbach & Brot, 1991; Ferla & Patrick, 2014; Brewster et al., 2021). This pathway is less parsimonious in terms of the number of steps and proteins involved and depends on sulfur to be first assimilated into cysteine (Seiflein & Lawrence, 2006; Hwang et al., 2007; Ferla & Patrick, 2014). To control the methionine biosynthesis pathway in *E. coli*, the enzymes in the trans-sulfurylation pathway are strictly regulated and feedback-inhibited by methionine and SAM (Ferla & Patrick, 2014; Sbodio et al., 2019). Thus, while *E. coli* is a valuable workhorse in synthetic biology, the trans-sulfuration pathway might be a limiting step when using it to bio-produce methionine. An alternative approach to bypass the inherent regulation of *E. coli* on the biosynthesis of methionine involves introducing genes from various organisms that are less prone to inhibition. Indeed, it was shown that MetX from *Leptospira meyeri* is not feedback-inhibited by methionine or S-adenosylmethionine (Bourhy et al., 1997). Although previous studies have demonstrated that modifying the methionine biosynthetic pathway of *E. coli* can effectively increase methionine levels, to date it had not been investigated whether simultaneously deleting both the *metA* and *metB* genes and replacing them with *metX* and *metY* to enable a full conversion from trans-to direct-sulfurylation would have a similar effect.

The primary objective of this study was to investigate the effect of replacing the trans-sulfurylation pathway in *E. coli* with the direct-sulfurylation machinery on methionine levels. Our findings demonstrate that the deletion of the *metA/B* genes in *E. coli* MG1655 resulted in a methionine auxotroph that could be complemented by the insertion of met*X/Y* genes from various sources. Furthermore, we found that the origin of the genes and their catalytic activity were closely associated with the ability of *E. coli* to produce methionine, leading to a significant increase in intra- and extra-cellular methionine levels.

## 2. Materials and Methods

### 2.1 Bacterial strains and growth conditions

The *E. coli* strain MG1655 is referred to as the WT strain and was used in this study for all genetic manipulation. The bacteria were routinely grown in a lysogeny broth (LB) medium at 37°C. For screening of the genetic variants, bacteria were grown in liquid or solid (supplemented with 1.5% agar) MOPS medium (8.73 g/L MOPS, 0.71 g/L tricine, 0.51 g/L NH_4_Cl, 0.05 g/L K_2_SO_4_, 2.92 g/L NaCl, 2.8 mg/L FeSO_4_, 0.074 mg/L, 0.1 g/L MgCl_2_ 1 ml/L trace elements, 30 mg/L K_2_HPO_4_, 2 g/L glucose, pH 7.3).

### 2.2 Generation of methionine auxotrophic mutants

All primers used in this study are listed in the supplementary information (Table S1). Genes in MG1655 were deleted by the lambda red recombinase procedure (Datsenko & Wanner, 2000), with the pkD4 plasmid carrying the Kn^R^ cassette serving as a template for PCR reactions. Mutations were verified using nearby locus-specific primers (Table S1), with the respective primers k2 or kt. Afterwards, the cassette was removed, and double/triple knockouts were further generated using a similar approach. Knockouts were generated in the following order Δ*met*A→ Δ*met*B→ Δ*metJ*, to form the bacterial strains Δ*met*A, Δ*met*AB and Δ*met*ABJ.

### 2.3 Complementation with a plasmid carrying the *metX/Y* genes

Electrocompetent Δ*metAB*/Δ*metABJ* mutants were transformed with the pCCI plasmid carrying the *metXY* synthetic operon. Following transformation, several colonies growing on LB agar plates supplemented with 30 µg/ml chloramphenicol were tested for the presence of the correct plasmid by colony PCR, with M13F and M13R primers. Positive clones were further screened for their ability to grow in methionine-depleted minimal media (MOPS). Briefly, cultures were prepared by inoculating a 5 ml MOPS medium with a single colony grown on LB plates and incubating it overnight at 37°C (constant orbital shaking 200 rpm). The culture was diluted 1,000-fold in a fresh MOPS medium, and 200 μl were then placed in each well of a 96-well plate (Costar). Bacteria were grown for 20 h, at 37°C (constant orbital shaking 280 rpm), and OD_600_ was measured every 16 min, using an Infinite M200 Plate Reader (Tecan). Each sample was tested in triplicates. For control, the MOPS medium was supplemented with 5-50 μg/ml methionine (Merck) or 0.1-0.5 mg/ml norleucine (Merck). Alternatively, a single colony grown on the LB medium was spread on MOPS agar plates, and growth following 24-72 h incubation at 37°C was visually inspected.

### 2.4 Construction of a *yjeH* overexpression plasmid

The *yjeH* gene was amplified from genomic DNA extracted from *E. coli* MG1655 using a forward primer that adds an *Nco*I restriction site (GCG<u>CCATGG</u>ATGAGTGGACTCAAACAAGAAC) and a reverse primer adding an *Xho*I restriction site (GCG<u>CTCGAG</u>TTATGTGGTTATGCCATTTTCC). The purified PCR product was digested with *Nco*I and *Xho*I and inserted into pTrcHis-a digested with the same enzymes. The correct construct was validated by sequencing.

### 2.5 Qualitative evaluation of extracellular methionine levels

For qualitative analysis of the methionine concentration in the medium, WT and methionine producing mutants were grown overnight at 37°C in a 5 ml MOPS medium and filtered through a 0.22 μm membrane to remove bacterial cells. The filtered cell-free medium was diluted two-fold in a fresh MOPS medium containing no methionine, ensuring all methionine in the new medium was secreted by the original bacteria. The new medium was tested for its ability to support the growth of a Δ*metAB* mutant in a minimal MOPS medium. Growth rate was determined by OD_600_ measurement.

### 2.6 Methionine extraction from lysate and medium to evaluate intra- and extra-cellular methionine levels

To evaluate the intracellular level of methionine, amino acids were extracted from cell pellets after centrifugation of 1 ml bacterial culture, using methanol:water:chloroform at a ratio of 1:1:2.5. After centrifugation, the crude extract was separated into polar and nonpolar phases by adding 300 µl water and 300 µl chloroform and centrifuging for 10 min. A 400 µl sample from the top polar phase were vacuum-dried. To evaluate extracellular methionine levels, amino acids were extracted from 500 µl of the bacterial culture medium by adding 500 µl chloroform and centrifuging for 10 min. A 400 µl sample from the upper polar phase was vacuum-dried. The latter fraction was dissolved in 40 µl of 20 mg/ml of methoxyamine hydrochloride in pyridine and incubated at 37°C for 2 h with vigorous shaking, followed by derivatization for 30 min in N-methyl-N(trimethylsilyl)-trifluoroacetamide at 37°C. One µl from each sample was used for methionine-level analysis, using GC-MS.

### 2.7 Evaluation of methionine levels by GC-MS

GC-MS analyses were carried out on Agilent 7890A GC-MS coupled with a mass selective detector (Agilent 5975c), a Gerstel multipurpose sampler (MPS2), and equipped with a BP5MS capillary column (SEG; 30 m, 0.25-mm i.d., and 0.25-mm thick). For free amino acid detection, the single-ion mass method was used. Amino acid standards of 5,10, 25, 50, 100 and 200 µM were used to generate standard calibration curves, and ribitol (2 mg/ml in HPLC-grade water) was used as an internal standard. Peak areas were calculated from the standard calibration curves and normalized to the ribitol signal.

### 2.8 Bacterial strains and plasmids used

Table 1 shows the different engineered strains and the terminology used in this study.

**Table 1.** Bacterial strains used in the study.

| Name in the manuscript | Description |
| --- | --- |
| Wildtype, WT | *E. coli* MG1655 |
| $\Delta metAB$ | *E. coli* MG1655 with a deletion of the *metA* and *metB* genes |
| $\Delta metABJ$ | *E. coli* MG1655 with a deletion of the *metA*, *metB* and *metJ* genes |
| $\Delta metABJ$-Y | *E. coli* MG1655 with a deletion of the *metA*, *metB* and *metJ* genes overexpressing the methionine exporter YjeH |
| DG/CM/LI/CG | Complementation of the engineered bacteria by the pCCI plasmid expressing *metX* and *metY* of the noted bacterial strain |

## 3. Results

### 3.1 Engineering of methionine auxotroph *E. coli*

To explore the option of converting the *E. coli* methionine biosynthesis pathway from trans-to direct-sulfurylation, we deleted two essential genes in the methionine pathway of *E. coli, met*A and *metB*, encoding for the enzymes HST and CGS, respectively (Figure 1). This deletion generated a methionine auxotroph *E. coli* strain (Δ*metAB*). Figure 2 shows growth curves of Δ*metAB* in a MOPS minimal medium containing glucose and ammonium chloride as the sole carbon and nitrogen sources, respectively, and K_2_SO_4_ as the sulfur source. To test the effect of methionine on the growth rate, we supplemented varying concentrations of external methionine, as indicated in Figure 2. While the WT bacteria grew normally without supplementation of methionine, the Δ*metAB* auxotroph was unable to grow. However, Δ*metAB* growth was rescued with the addition of methionine. At a concentration of 25 µg/ml methionine, Δ*metAB* growth reached maximal levels and the cell density resembled that of the WT bacteria, demonstrating that methionine was indeed the limiting growth factor.

**Figure 2.**
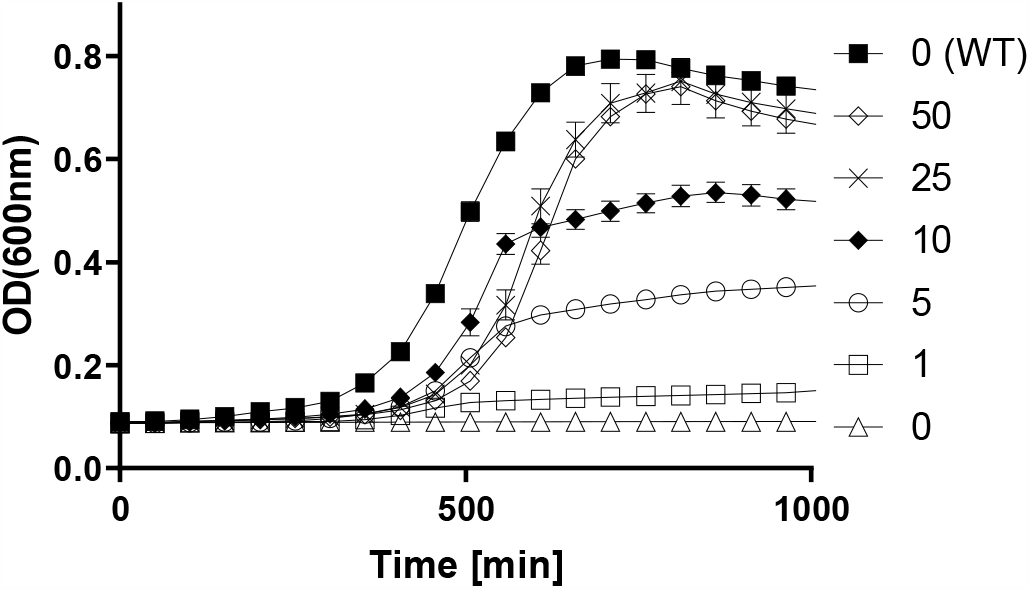
*E. coli* Δ*metAB* is auxotrophic for methionine. WT and Δ*metAB* were grown in a minimal MOPS medium with or without supplementation of external methionine for 900 min at 37°C. The legend shows methionine concentration in μg/ml

### 3.2 Complementation of *E. coli* Δ*met*AB by *metX* and *metY* genes from different bacterial genomes

With the aim of exploring *E. coli’s* ability to synthesize methionine via the direct-sulfurylation pathway using the Met*X* and Met*Y* enzymes, we cloned four different *metY/X* gene pairs to complement the Δ*metAB* auxotroph bacteria. The selection of *metY/X* gene pairs was based on a previously characterized dataset of MetX enzymes from various bacteria (Bastard et al., 2017). We applied two criteria to select the strains. First, the MetX enzymes should exhibit a range of catalytic activity between 10^3^-10^4^ nmol·min^-1^·mg^-1^ (of *O*-acetyl L-homoserine formation). Second, the relevant bacterial genome should contain a sequence for the counterpart *metY* gene adjacent to the *metX* gene, indicating a mini operon of two genes with coordinated expression and function.

Based on the above rationale, we selected *metY*/*metX* pairs from four bacterial strains: i) *Corynebacterium glutamicum* (CG), ii) *Leptospira interrogans* (LI), iii) *Cyclobacterium marinum* (CM) and iv) *Deinococcus geothermalis* (DG). Table 2 summarizes the UniProt entry identifiers of each protein. Sequence identity between the various MetX and MetY proteins is provided in table S2 of the supplementary information.

**Table 2.** UniProt identifiers of the enzymes encoded by the genes used to complement the auxotroph bacteria.

| Bacterial strain | metX UniProt ID | metY UniProt ID |
| --- | --- | --- |
| *Leptospira interrogans* | Q8F4I0 | P94890 |
| *Corynebacterium glutamicum* | O68640 | Q79VI4 |
| *Deinococcus geothermalis* | Q1J115 | Q1J114 |
| *Cyclobacterium marinum* | G0J5N4 | G0J5N3 |

To design a uniform construct for expression in *E. coli*, the *metX* and *metY* genes were codon-optimized and cloned into a pCCl plasmid (Wild et al., 2002; Venkova-Canova et al., 2004) together with a synthetic constitutive promoter, a synthetic ribosome binding site (RBS), and a synthetic terminator. The regulatory elements ensured constitutive expression, and the use of the pCCI plasmid enabled the maintenance of the inserted genes at low to a single-copy number, similar to the genomic copy number of the corresponding genes. Figure 3A shows a schematic illustration of the constructed operon. The complete gene sequences and their accession numbers are depicted in the supplementary information. Following the transformation of the auxotroph bacteria with the plasmids, we evaluated the ability of the complemented Δ*metAB* strain to grow in a liquid minimal MOPS medium. As shown in Figure 3B, the Δ*metAB* bacteria complemented with *metY/X* from DG and CM (Δ*metAB*-DG and Δ*metAB*-CM, respectively) successfully complemented the auxotroph bacteria and reached a similar growth rate and final cell density as those of the WT after ∼800 min. The complemented bacteria carrying *metY*/*X* of LI and CG (Δ*metAB*-LI and Δ*metAB*-CG in Figure 3B) did not grow under these conditions. Similar growth patterns were observed on minimal medium agar plates (Figure 3C), indicating that the CM- and DG-complemented strains were able to produce methionine at levels sufficient to maintain their growth.

**Figure 3.**
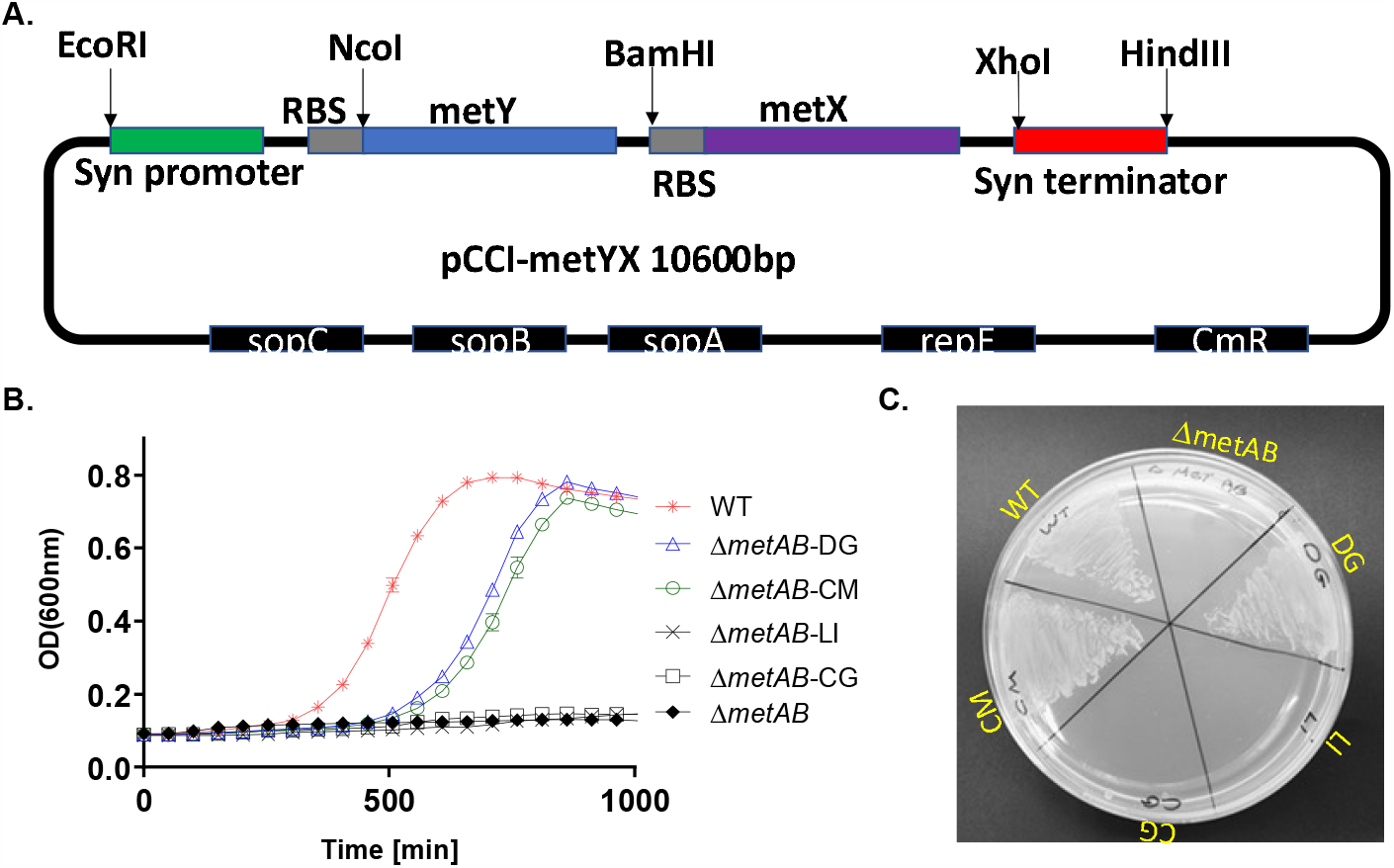
Complementation of Δ*metAB* with *metX/Y* gene pairs. **A**. Schematic illustration of the synthetic *metYX* operon on a low-copy plasmid used to complement Δ*metAB*. A synthetic operon consisting of the *metY* and *metX* genes was constructed by adding a synthetic constitutive promoter, a ribosome binding site (RBS) for each gene, and a synthetic terminator. Restriction sites were included to facilitate rearrangement and analysis of mutant genes. **B**. Growth curves of the complemented Δ*metAB* strains on a minimal MOPS medium. WT: wild type MG1655; Δ*metAB*: WT with deletion of *metAB* genes; Δ*metAB*-DG/CM/LI/CG: Δ*metAB* complemented with a pCCl plasmid expressing *metX* and *metY* of the indicated bacterial strain. **C**. Growth of the complemented Δ*metAB* strains on a MOPS minimal-medium agar plate incubated at 37°C for 24 h.

To quantify intracellular and extracellular methionine levels in the complemented bacteria, the methionine levels were evaluated using GC-MS and compared to those of the WT bacteria. The Δ*metAB*-DG *and* Δ*metAB*-CM strains exhibited a five-fold enhancement of intracellular methionine levels compared to WT (Figure 4A). Analysis of the extracellular methionine in the growth medium indicated that Δ*metAB*-DG exhibited significantly enhanced accumulation of extracellular methionine as compared to WT (18 fold), Although the difference in methionine accumulation was not significant the Δ*metAB*-CM also showed a five-fold increase compared to the WT (Figure 4B).

**Figure 4.**
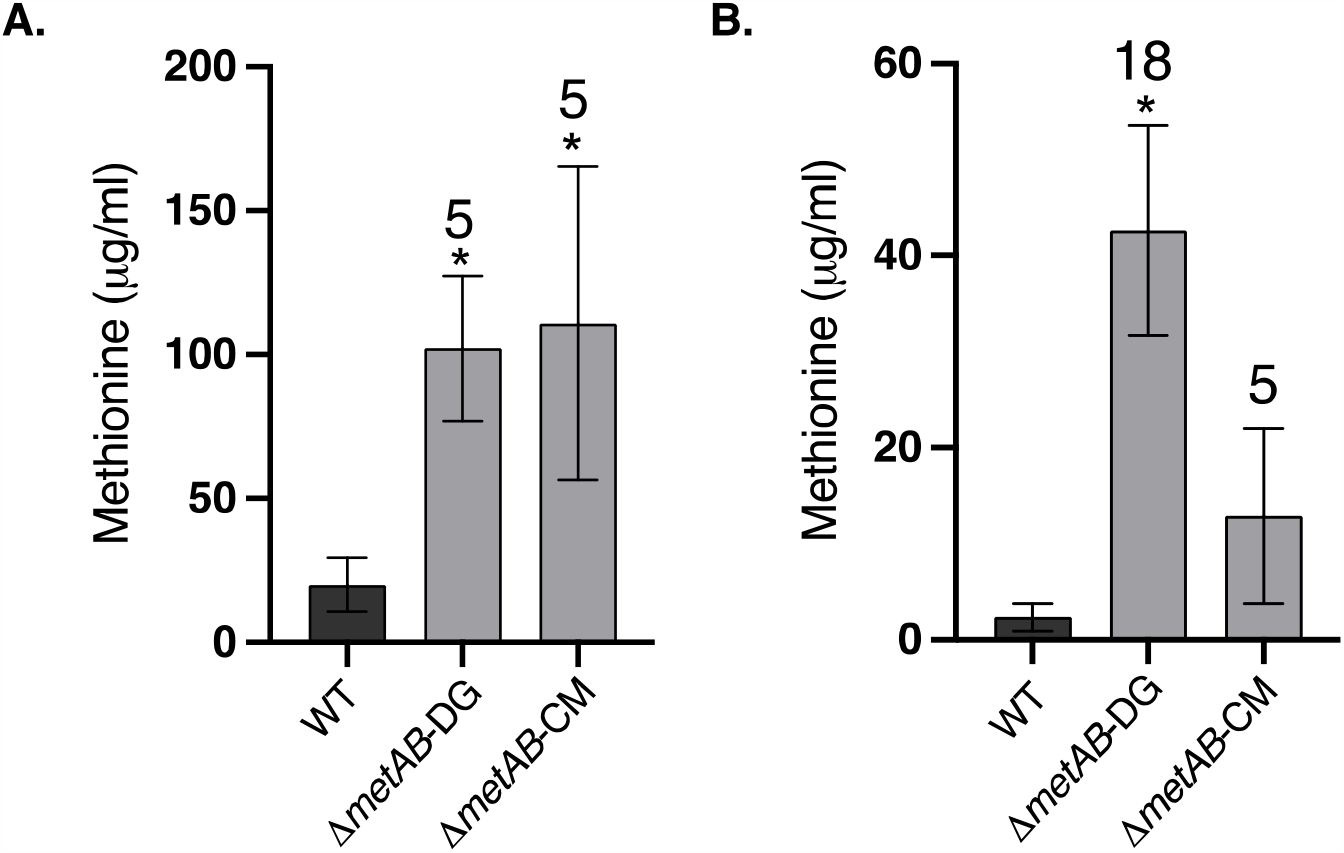
Biosynthesis of methionine by *E. coli* Δ*metAB* complemented with *metY/X* pairs. WT and complemented *E. coli* Δ*metAB* were grown in a minimal MOPS medium at 37°C for 24 h, after which the cells were separated from the growth medium. The amount of methionine in each fraction was evaluated using GC-MS. **A**. Intracellular methionine accumulated by WT *E. coli*, Δ*metAB*-DG and Δ*metAB*-CM, reported as μg/ml. **B**. Extra-cellular methionine accumulated in the growth media by WT *E. coli*, Δ*metAB*-DG and Δ*metAB*-CM, reported as μg/ml. The results are presented as means ± SD of three to four replicates for each sample. Significance between WT and the different bacterial strain was calculated according to the Student’s t-test (P < 0.05) and is identified by an asterisk. The numbers on top of the bars indicate the fold increase relative to the WT in each panel.

### 3.3 Evaluation of intra- and extracellular methionine levels of *E. coli* with a deletion of *metJ* and overexpression of YjeH

After establishing *E. coli* strains expressing the *metY*/*X* of DG and CM in a Δ*metAB* background, we further explored the effect of additional genetic variations related to the methionine biosynthetic pathway. More specifically, we deleted the gene encoding for the MetJ repressor that is known to strictly repress the transcription of multiple genes in the *E. coli* methionine biosynthetic pathway (*metA, metB, metC, metE/H*) in response to elevated methionine levels (Saint-Girons et al., 1984; Augustus et al., 2010).

To evaluate the ability of the engineered strains to produce methionine, the bacteria were cultured in a minimal medium until reaching OD_600_=2.5. The intracellular and extracellular levels of methionine were evaluated using GC-MS (Figure 5). Deletion of *metJ* on a WT background led to a 9-fold enhancement in the level of intracellular methionine relative to WT (Figure 5A), while no change was detected in the extracellular methionine levels (Figure 5B). Deletion of *metJ* on the Δ*metAB*-CM background led to a 16- and 45-fold increase in intracellular and extracellular methionine levels, respectively, while in Δ*metABJ*-DG, the levels increased by 11- and 95-fold, respectively, relative to WT (Figure 5A-B).

**Figure 5.**
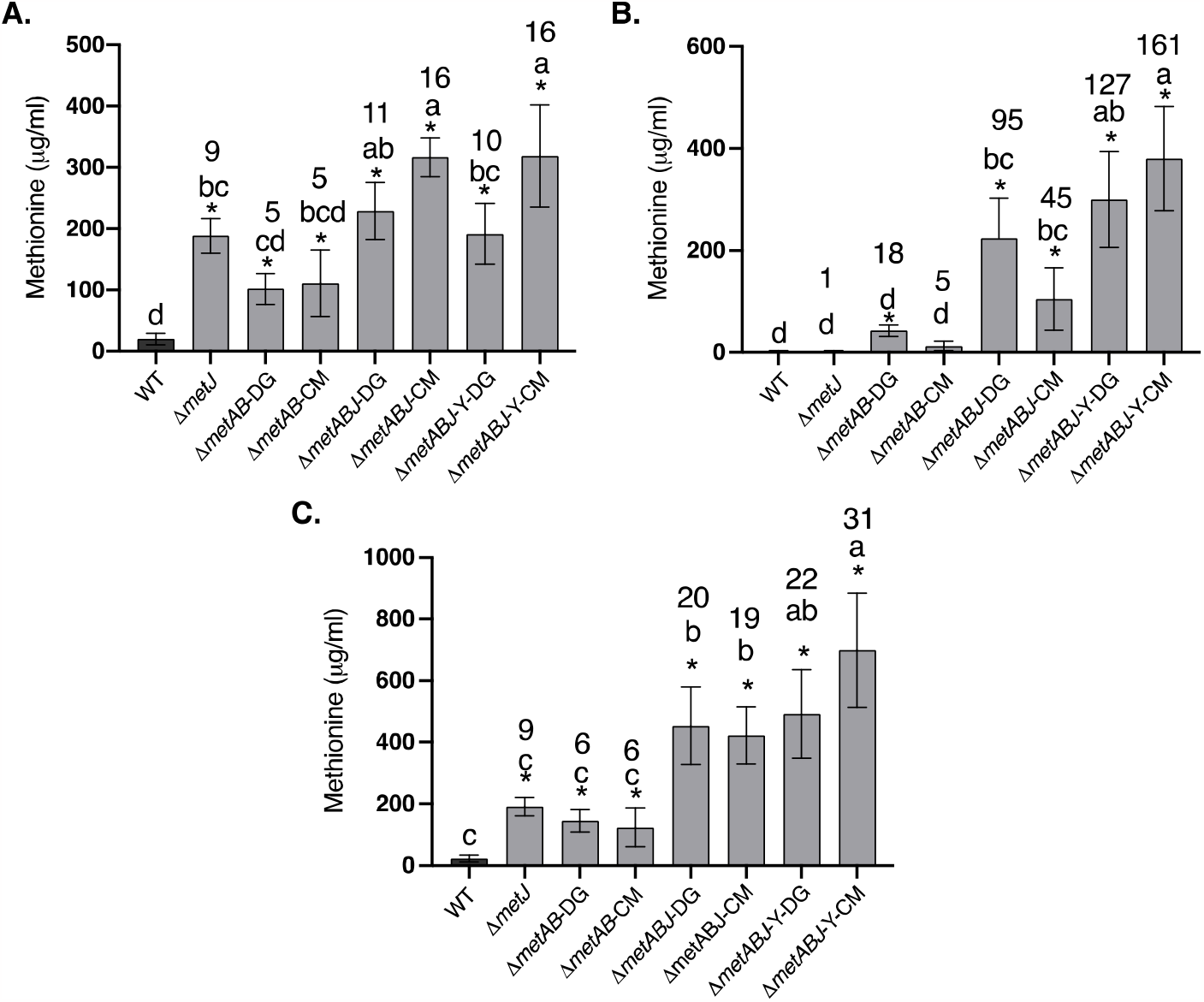
Production of methionine by Δ*metABJ* overexpressing YjeH and complemented by *metY/X* from CM or DG. Comparison of: **A**. intracellular; **B**. extracellular; and **C**. total methionine levels that were quantified by GC-MS. Peak areas were normalized to norleucine internal control, and total methionine levels were calculated according to the standard calibration curves. The results are presented as means ± SD of three or four replicates for each sample. Significance between bacterial strains was calculated according to the Turkey-Kramer HSD test (p < 0.05) and is identified by different small letters. Significance between WT and the different bacterial strains was calculated according to the Student’s t-test (P < 0.05) and is identified by an asterisk. The numbers on top of the bars in each panel indicate the fold-increase relative to the WT in each panel.

To further increase methionine levels in the growth medium and reduce the level of inhibition on methionine-feedback sensitive enzymes or regulators, we also targeted the *E. coli* methionine exporter protein YjeH. This exporter was shown to have a strong positive effect on extracellular methionine accumulation and to reduce the methionine content inside the bacterial cells (Liu et al., 2015). Therefore, its gene was cloned to facilitate overexpression and to enable enhanced methionine efflux to the medium. The *metY/X* plasmid carrying *metY/X* from DG or CM was than introduced into Δ*metABJ*-Y, resulting in Δ*metABJ*-Y-CM and Δ*metABJ*-Y-DG. This procedure led to similar intracellular methionine levels as in Δ*metAB*-CM and Δ*metABJ*-DG, but it significantly increased the levels of extracellular methionine by 161- and 127-fold, respectively, relative to WT (Figure 5B). Overall, methionine levels (combining the intra and extracellular methionine) increased by up to 31-fold over the WT and reached up to 700 μg/ml (Figure 5C).

### 3.4 Bioavailability of extracellular methionine secreted from the engineered Δ*metABJ-Y*-DG and Δ*metABJ-Y*-CM strains and its potential use as a methionine supplement

To confirm the bioavailability of the extracellular methionine secreted from each engineered strain using an orthogonal approach, we collected the spent medium at the end of the bacterial growth phase of each strain and filtered it through a 0.22 μm membrane. The filtered spent medium was then added to a fresh methionine-free minimal MOPS medium at a 1:1 ratio. Thus, methionine could only be delivered from the filtered spent medium (Figure 6A). The mixed medium was then evaluated for its ability to support the growth of Δ*metAB* auxotroph. Figure 6 shows the growth curves of the Δ*metAB* auxotroph bacteria grown in the mixed medium originated from the spent medium of the DG-(Figure 6B) and CM-(Figure 6C) complemented strains. The highest cell density was observed when the Δ*metAB* auxotroph was cultured with spent medium originating from Δ*metABJ*-Y-CM/DG, suggesting that this strain exported the highest methionine levels. On the other hand, the Δ*metAB* auxotroph did not grow with medium originated from the WT, indicating for the lack of methionine in its medium. Both results are congruent with the methionine levels that were measured for these strains using GC-MS (Figure 5).

**Figure 6.**
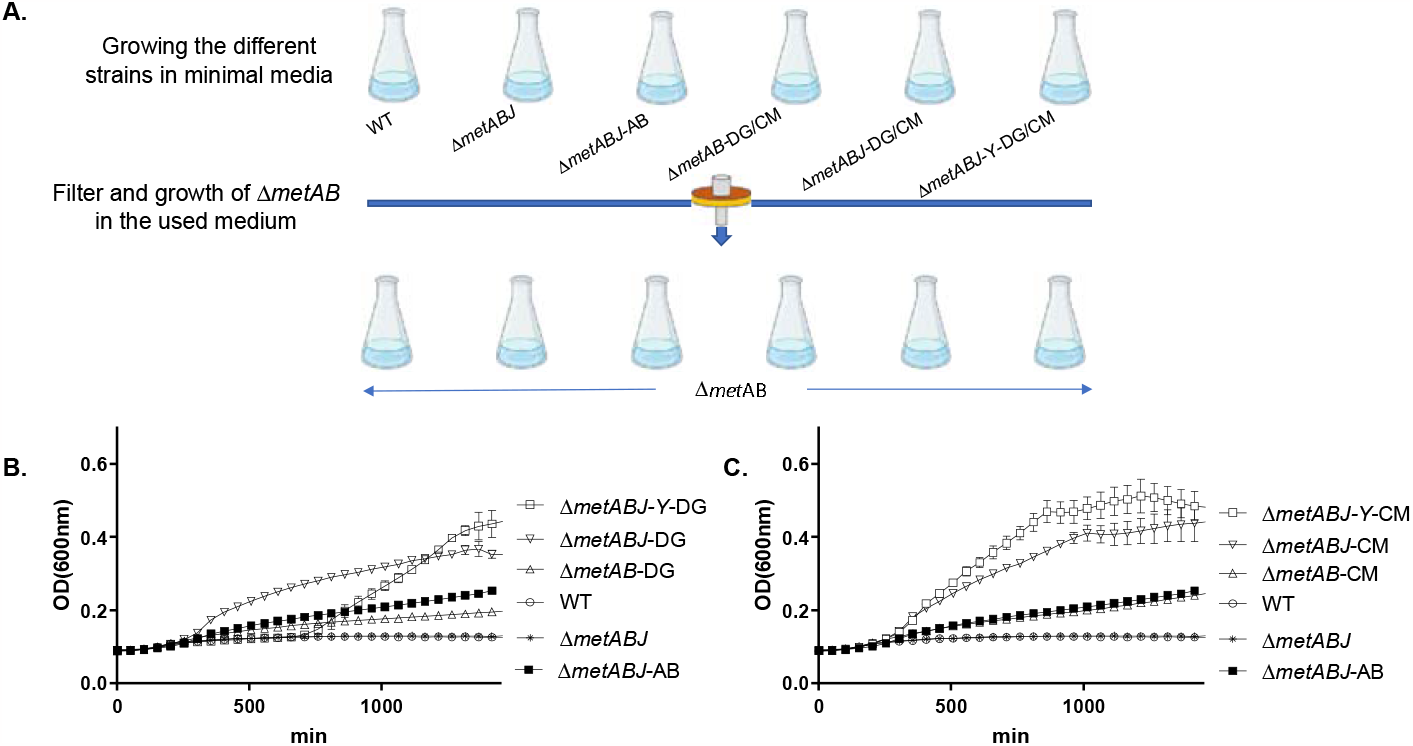
Growth curves of methionine-auxotroph *E. coli* in spent medium of each strain. **A**. Illustration of the experimental scheme used to evaluate methionine level in the medium following the growth of each strain. **B**. Growth curves in medium from DG strains. **C**. Growth curves in medium from CM strains. All curves show the growth of the auxotroph *E. coli* Δ*metAB* in fresh MOPS minimal medium supplemented with spent medium filtered following the growth of the indicated strains.

## Discussion

*E. coli* is currently the most extensively explored organism for enhancing the bioproduction of natural amino acids, including L-methionine (Sanchez et al., 2017; Wendisch, 2020; Mohany et al., 2021; Niu et al., 2023; Shen et al., 2023). While most studies involving metabolic engineering for increased methionine production in *E. coli* were conducted in *E. coli* W3110 and reported methionine levels of up to 18 g/L using medium supplemented with yeast extract and vitamins (Zhou, et al., 2019, Niu, et al., 2023), in the present study we used MOPS minimal medium and utilized the close relative, *E. coli* MG1655 (as per Huang et al., 2017; Li et al., 2017), which is used interchangeably and was proposed to be a good producer of bioproducts (Hayashi et al., 2006; Khankal et al., 2008; Monk et al., 2016). Regardless of the bacterial strain, efforts to enhance methionine levels in *E. coli* mostly involve the engineering of multifaceted cellular pathways that aim to release negative feedback regulation alongside optimizing the utilization of methionine precursors. While this strategy has resulted in significant improvements, it relies on harnessing the natural trans-sulfurylation pathway of *E. coli*. Methionine biosynthesis by the direct-sulfurylation pathway is much more abundant in the bacterial kingdom. However, it has been characterized in a relatively limited number of strains (Ferla & Patrick, 2014). Moreover, only limited data is available on the catalytic properties of the central MetY enzymes (Hwang et al., 2007; Kulikova et al., 2019; Kulikova et al., 2021) and their related 3D structures (Imagawa et al., 2005, 2007; Tran et al., 2011; Brewster et al., 2021).

Enzymes of the direct-sulfurylation pathways are versatile and can process various substrates (Foglino et al., 1995; Hacham et al., 2003). Indeed, previous studies showed that *E. coli* can grow with such enzymes (Bourhy et al., 1997; Schipp et al., 2020). Thus, to reduce the high energy cost and tight regulation of trans-sulfurylation, the current study aimed to replace the natural enzymes in *E. coli* with their counterpart from the direct-sulfurylation pathway. To that end, we explored the ability to complement the auxotroph Δ*metAB* strain with *metX* and *metY* enzymes from various bacteria completely forming direct-sulfurylation within the *E. coli*. The heart of the effort involved replacing the enzymes HST and CgS with HAT and OAHS. We inserted the genes encoding MetX (HAT) and MetY (OAHS) into the methionine auxotroph strain via a plasmid containing a synthetic mini operon of *metY* followed by *metX* (Figure 2A). The transformation of the auxotrophic strain with a plasmid carrying the *metY/X* genes from DG and CM allowed for bacterial growth without the external addition of methionine. Notably, the insertion of *metX* and *metY* from CG and LI failed to complement the auxotroph *E. coli*. This could be due to these enzymes’ reduced efficiency compared to the enzymes of DG and CM. However, it is possible that the activity of these enzymes requires additional co-factors and/or certain conditions that are not present in the context of the trans-sulfurylation pathway within *E. coli*. An additional explanation could be that a reduced expression level or faster degradation contributed to the inability of the bacteria to grow. Regardless of the exact mechanism, this finding suggests that large variability exists in the activity of the different enzymes when complemented into *E. coli*. Therefore, the screening of additional genes from multiple organisms may further benefit methionine accumulation.

Several mechanisms could explain the higher methionine accumulation in the strains complemented with enzymes from the DG and CM strains relative to the WT containing the *met*A and *met*B genes on the same construct (Figure 6B-C). In the latter, methionine biosynthesis is under strict regulation, including on the protein level (Usuda & Kurahashi, 2005; Huang et al., 2017). Indeed, deletion of *metJ* resulted in higher methionine biosynthesis in the WT strain, showing that the release of regulation at the transcription level is an important factor for enhancing methionine biosynthesis (Usuda & Kurahashi, 2005; Li et al., 2017). Without the transcriptional regulation in the Δ*metJ* strain, deletion of *metA* and *metB* together with complementation with *metX* and *metY* (Δ*metABJ*-DG/CM) further pushed the levels of methionine above those found in Δ*metAB*-DG/CM. This finding could be due to the higher rate of enzymatic activity of *metX* in comparison to the rate-limiting enzyme *metA* in the trans-sulfurylation pathway (Bastard et al., 2017) in addition to the reduced regulation of other important genes in the pathway, such as *metE* and *metH* (Figure 1). Indeed, it was previously shown that MetX from LI expressed in *E. coli* was not affected by feedback inhibition imparted by high levels of methionine or SAM (Bourhy et al., 1997). Of note, the higher levels of methionine observed in the Δ*metABJ*-Y-CM compared to Δ*metABJ*-CM suggest that excess of methionine inside the cells is controlled by other factors, some of which are yet unknown. Indeed, when the YjeH transporter was overexpressed, it enhanced the cells efflux and enabled the bacteria to produce more methionine.

The enhancement of methionine biosynthesis in the engineered *E. coli* warrants screening of additional MetX and MetY enzymes of other strains, to further characterize their ability to support methionine production. In addition, it may be possible to boost methionine levels by optimizing growth conditions with alternative sulfur sources and introducing additional modifications to the direct-sulfurylation pathway that aim to enhance metabolic flux and methionine export. Discovery of additional factors and their subsequent genetic alteration may further increase levels of methionine. These alterations can be achieved by classical strain improvement, using inhibitors such as norleucine, or by building new genetic circuits to control the expression of relevant genes. Our results show that the MetAB enzymes could be a limiting step in methionine biosynthesis regardless of additional modifications applied to the cell, and that the use of direct-sulfurylation MetYX enzymes dramatically enhanced methionine production (Figure 5).

Additionally, our findings demonstrate that through the substitution of trans-sulfurylation with direct-sulfurylation, elevated levels of methionine can be exported and accumulated in the growth medium. This bioavailable methionine successfully supported the growth of the Δ*metAB* auxotroph bacteria and thus has promising applications in fields such as animal feed and mammalian cell culture cultivation. As such, these findings pave the way for further advancements in the utilization of bacterial methionine.

## Conclusions

Harnessing *E. coli* to produce L-methionine presents a promising avenue for enhanced production; however, it necessitates the modification of numerous regulatory and enzymatic bottlenecks throughout the biosynthetic pathway. Our findings suggest that by substituting the trans-sulfurylation *metA* and *metB* genes with the direct-sulfurylation *metX* and *metY* genes, methionine production in minimal medium can be significantly enhanced up to 700 mg/L.

## Supporting information

supplementary tables and sequences

## Acknowledgments

This research was supported by the Israeli Ministry of Agriculture (grant# 21-36-0003).

## Author contributions

N.G., Y.H., H.H., I.S., and M.G. performed the experimental research. Y.H. performed all mass-spec experiments. I.B. performed the bioinformatic analysis. N.G., Y.H., R.A., I.Y., and M.G. analyzed the data. Y.H., R.A., I.Y., and M.G. conceived of the research and wrote the manuscript. All authors reviewed and approved the manuscript.

