## supplementary tables and sequences for "Conversion of methionine biosynthesis in *E. coli* from trans- to direct-sulfurylation enhances extracellular methionine levels"

**supplementary information**

**TABLES**

Table S1. Primers used in this study.

| Primer name | Primer sequence | Description |
| --- | --- | --- |
| **MetA-D-For** | **CGAGCTACCCGCCGTCAATTTCTTGCGTGAAGAAAACGTCTTTGTGATGA**CATATGAATATCCTCCTTAG | Generation of Δ*met*A mutation |
| **MetA-D-Rev** | **TGTGCCGTAGATCGTATGGCGTGATCTGGTAGACGTAATAGTTGAGCCAG**TGTAGGCTGGAGCTGCTTCG | Generation of Δ*met*A mutation |
| **MetB-D-For** | ACAGGCCACCATCGCAGTGCGTAGCGGGTTAAATGACGACGAACAGTATGCATATGAATATCCTCCTTAG | Generation of Δ*met*B mutation |
| **MetB-D-Rev** | CGGAAGCCATTTTCCAGGTCGGCAATTAAATCTTCGCCATCTTCAATACCTGTAGGCTGGAGCTGCTTCG | Generation of Δ*met*B mutation |
| **MetJ-D-For** | **GGAGCGGCGAATATATCAGCCCATACGCTGAGCACGGCAAGAAGAGTGAA**TGTAGGCTGGAGCTGCTTCG | Generation of Δ*met*J mutation |
| **MetJ-D-Rev** | **CACGTCTCCGGGTTAATCCCCATCTCACGCATGATCTCTTTTGCCGCTTC**CATATGAATATCCTCCTTAG | Generation of Δ*met*J mutation |
| **MetA-For** | GTTATGCCGATTCGTGTGCC | Verification of Δ*met*A mutation |
| **MetA-Rev** | AGGTAAGGTGCTGAATCGCT | Verification of Δ*met*A mutation |
| **MetB-For** | ACATTTCACCGACAAAGCCC | Verification of Δ*met*B mutation |
| **MetB-Rev** | TTTACCCCTTGTTTGCAGCC | Verification of Δ*met*B mutation |
| **MetJ-For** | ACTGTGTGGTCTGGTCTCAA | Verification of Δ*met*J mutation |
| **MetJ-Rev** | CTGATAAGCGTAGCGCATCA | Verification of Δ*met*J mutation |
| **yjeH-For** | GCGCCATGGATGAGTGGACTCAAACAAGAAC | Amplification of genomic copy of yjeH and addition of a NcoI site for cloning (underlined) |
| **yjeH-Rev** | GCGCTCGAGTTATGTGGTTATGCCATTTTCC | Amplification of genomic copy of yjeH and addition of a XhoI site for cloning (underlined) |

- Underlined sequences refer to sequences derived from pKD4 plasmid (Datsenko & Wanner, 2000)

**Sequence identity**

Table S2 shows the identity between the four selected MetX proteins (33.5-37%) and between the four selected MetY proteins (43.9-56.7%).

Table S2. Multiple sequence alignment of the selected MetX and MetY sequences.

|  | ***MetX_CG*** | ***MetX_LI*** | ***MetX_CM*** | ***MetX_DG*** |
| --- | --- | --- | --- | --- |
| ***MetX_CG*** | *100* | *35.92* | *36.95* | *33.54* |
| ***MetX_LI*** |  | *100* | *37.21* | *35.31* |
| ***MetX_CM*** |  |  | *100* | *36.96* |
| ***MetX_DG*** |  |  |  | *100* |

|  | ***metY_CG*** | ***metY_LI*** | ***metY_CM*** | ***metY_DG*** |
| --- | --- | --- | --- | --- |
| ***MetY_CG*** | *100* | *49.76* | *43.87* | *47.18* |
| ***MetY_LI*** |  | *100* | *55.2* | *51.96* |
| ***MetY_CM*** |  |  | *100* | *56.68* |
| ***MetY_DG*** |  |  |  | *100* |

*Alignment was conducted using MUSCLE and the numbers represent percent identity*

Supplementary text 1. Construct sequences

> metYX_CG_ OQ291222

gaattcTTTATTCTTGACACTAGTCGGCCAAAATGATATAATACCTGAGTTTAACTTTAAGAGAGGTATATATTACCATGGGTCCGAAGTACGACAACAGCAACGCGGATCAGTGGGGCTTCGAGACCCGTAGCATCCACGCGGGTCAGAGCGTTGACGCGCAAACCAGCGCGCGTAACCTGCCGATTTACCAGAGCACCGCGTTCGTGTTTGACAGCGCGGAGCACGCGAAACAACGTTTCGCGCTGGAAGATCTGGGCCCGGTTTATAGCCGTCTGACCAACCCGACCGTGGAGGCGCTGGAAAACCGTATTGCGAGCCTGGAGGGTGGCGTTCATGCGGTGGCGTTTAGCAGCGGTCAGGCGGCGACCACCAACGCGATCCTGAACCTGGCGGGTGCGGGTGACCACATTGTTACCAGCCCGCGTCTGTATGGTGGCACCGAAACCCTGTTCCTGATCACCCTGAACCGTCTGGGCATTGATGTTAGCTTTGTGGAGAACCCGGATGATCCGGAAAGCTGGCAGGCGGCGGTTCAACCGAACACCAAGGCGTTCTTTGGCGAGACCTTTGCGAACCCGCAAGCGGACGTGCTGGATATCCCGGCGGTTGCGGAAGTGGCGCACCGTAACAGCGTTCCGCTGATCATTGACAACACCATTGCGACCGCGGCGCTGGTGCGTCCGCTGGAGCTGGGTGCGGATGTGGTTGTGGCGAGCCTGACCAAGTTCTACACCGGTAACGGCAGCGGTCTGGGTGGCGTTCTGATCGACGGTGGCAAATTTGATTGGACCGTGGAAAAGGACGGCAAAAGCGTTTTCCCGTATTTTGTTACCCCGGATGCGGCGTACCACGGTCTGAAGTATGCGGATCTGGGTGCGCCGGCGTTTGGTCTGAAAGTTCGTGTGGGCCTGCTGCGTGACACCGGTAGCACCCTGAGCGCGTTTAACGCGTGGGCGGCGGTTCAAGGCATCGATACCCTGAGCCTGCGTCTGGAGCGTCACAACGAAAACGCGATTAAGGTGGCGGAGTTCCTGAACAACCACGAGAAGGTTGAAAAAGTGAACTTTGCGGGTCTGAAGGATAGCCCGTGGTACGCGACCAAGGAAAAACTGGGCCTGAAATATACCGGTAGCGTGCTGACCTTCGAGATCAAGGGTGGCAAAGACGAAGCGTGGGCGTTTATTGATGCGCTGAAACTGCACAGCAACCTGGCGAACATCGGCGACGTTCGTAGCCTGGTTGTGCATCCGGCGACCACCACCCATAGCCAAAGCGATGAGGCGGGCCTGGCGCGTGCGGGTGTGACCCAAAGCACCGTTCGTCTGAGCGTGGGTATCGAGACCATTGACGATATCATTGCGGACCTGGAAGGTGGCTTCGCGGCGATTTAAGGATCCAGAGGTATATATTAatgCCGACCCTGGCGCCGAGCGGTCAGCTGGAGATCCAAGCGATTGGTGACGTTAGCACCGAGGCGGGCGCGATCATTACCAACGCGGAAATTGCGTACCACCGTTGGGGTGAGTATCGTGTGGACAAAGAAGGCCGTAGCAACGTGGTTCTGATCGAACACGCGCTGACCGGTGATAGCAACGCGGCGGACTGGTGGGCGGATCTGCTGGGTCCGGGCAAGGCGATCAACACCGACATTTACTGCGTTATCTGCACCAACGTGATCGGTGGCTGCAACGGCAGCACCGGTCCGGGCAGCATGCACCCGGATGGTAACTTCTGGGGCAACCGTTTTCCGGCGACCAGCATTCGTGACCAGGTTAACGCGGAGAAACAATTCCTGGATGCGCTGGGTATTACCACCGTTGCGGCGGTGCTGGGTGGCAGCATGGGTGGCGCGCGTACCCTGGAGTGGGCGGCGATGTATCCGGAAACCGTTGGTGCGGCGGCGGTGCTGGCGGTTAGCGCGCGTGCGAGCGCGTGGCAGATCGGCATTCAGAGCGCGCAAATCAAGGCGATTGAAAACGATCACCACTGGCACGAGGGTAACTACTATGAAAGCGGCTGCAACCCGGCGACCGGTCTGGGTGCGGCGCGTCGTATTGCGCACCTGACCTACCGTGGTGAGCTGGAAATCGACGAGCGTTTTGGCACCAAGGCGCAGAAAAACGAAAACCCGCTGGGTCCGTATCGTAAGCCGGATCAACGTTTCGCGGTTGAGAGCTACCTGGACTATCAGGCGGATAAACTGGTTCAACGTTTTGACGCGGGTAGCTACGTGCTGCTGACCGATGCGCTGAACCGTCACGACATTGGCCGTGATCGTGGTGGCCTGAACAAGGCGCTGGAGAGCATTAAAGTGCCGGTTCTGGTGGCGGGCGTTGACACCGATATCCTGTACCCGTATCACCAGCAAGAACACCTGAGCCGTAACCTGGGTAACCTGCTGGCGATGGCGAAAATCGTTAGCCCGGTGGGTCATGATGCGTTCCTGACCGAAAGCCGTCAAATGGATCGTATTGTGCGTAACTTCTTTAGCCTGATCAGCCCGGACGAGGATAACCCGAGCACCTACATTGAATTTTATATCTAACTCGAGCAACCTGGAGGCGGGCGCAGGCCCGCCTTTTaagctt

> metYX_DG_OQ291223

gaattcTTTATTCTTGACACTAGTCGGCCAAAATGATATAATACCTGAGTTTAACTTTAAGAGAGGTATATATTAccATGGCGAGCAACACCCTGCACTTCGAGACCCTGCAAGTGCACGCGGGTCAACATCCGGACCCGGCGACCGGCGCGCAAGCGGTGCCGATTTACGCGACCAACGCGTATGTTTTTGAAAGCCCGGAACATGCGGCGGACCTGTTCGGTCTGCGTGCGTTTGGCAACATTTACAGCCGTATCATGAACCCGACCAACGCGGTTCTGGAGGAACGTATTGCGGCGCTGGAAGGTGGCGTGGGTGCGCTGGCGGTTGCGAGCGGTCATGCGGCGCAGTTCCTGGCGATTACCACCGTGGCGCAAGCGGGTGATAACATCGTTAGCACCCCGAACCTGTACGGTGGCACCGTGAACCAATTTCGTGTTACCCTGCGTCGTCTGGGCATTGAAGTGCGTTTCACCAGCAAGGACGAACGTCCGGAGGAATTTGCGGCGCTGATCGACGATCGTACCCGTGCGGTTTATCTGGAGACCCTGGGTAACCCGGCGCTGAACGTGCCGGACTTTGAGGGTATTGCGGAAGTTGCGCATGCGCGTGGCGTGGCGGTTTTTGTGGATAACACCTTTGGTGCGGGTGGCTACTATTGCCAACCGCTGCGTCACGGCGCGGATGTGGTTCTGCACAGCGCGAGCAAATGGATTGGTGGCCACGGTAACGGCATCGGTGGTCTGCTGGTTGACGGTGGCACCTTTGATTGGGGTAACGGCCGTTACCCGCTGCTGACCGAGCCGAGCCCGAGCTATCACGGTCTGAGCTTCTGGGAGGCGTTTGGTGAGGGTAACGCGCTGGGTCTGCCGAACATTGCGTTCATTACCCGTGCGCGTACCGAAGGTCTGCGTGATCTGGGTCCGACCCTGGCGCCGCAGCAAGCGTGGCAGTTTCTGCAAGGTGTGGAGACCCTGAGCCTGCGTGCGGAACGTCATGCGCAGAACGCGCTGGCGCTGGCGAGCTGGCTGAGCGGTCACCCGGATGTGAGCCGTGTTACCTATCCGGGCCTGAGCAACCACCCGCACTACGATCGTGCGCAAACCTATCTGCCGCGTGGTGCGGGTGCGGTTCTGACCTTTGAGCTGCGTGGTGGCCGTGCGGCGGGTGAAGCGTTTATTGGTGCGGTGCGTCTGGCGCAGCATGTGGCGAACGTTGGTGACACCCGTACCCTGGTTATTCATCCGGCGAGCACCACCCACAGCCAGCTGGATGAAGCGGCGCAAGCGGCGGCGGGTGTGACCCCGGGCCTGGTTCGTGTGAGCGTTGGTATCGAGCACATTGACGATATCCGTGAAGATTTTGCGCAGGCGCTGGCGACCGCGCTGGTTGATGCGGAGGGTGCGTAAggatccAGAGGTATATATTAATGCGTATCCAGCGTTACATTCTGATGACCGCGCTGATCAGCCAACCGGACCTGCTGCCGCCGCCGGCGCCGGAGCGTTGCCCGCCGCAGCAAACCGCGCGTCTGTTCCGTGAAACCCCGCTGCTGCTGGACTGCGGTCAGGTGGTTCAAGATGTGCGTGTTGCGTACCACACCTATGGCACCCCGAGCGACCATGCGATCCTGGTGCTGCATGCGCTGACCGGCACCAGCGCGGTTCATGAGTGGTGGCCGGATTTTCTGGGTGAAGGCAAGCCGCTGGACCCGACCCGTGATTATATTGTTTGCGCGAACGTTCTGGGTGGCTGCGCGGGTAGCACCGGTCCGGCGGAGCTGCCGCGTGTGAACGGTGAAGACCCGCCGCTGACCCTGCGTGATATGGCGCGTGTGGGTCGTGCGCTGCTGGAGGAACTGGGCGTTCGTCGTGTGAGCGTTATTGGTGCGAGCATGGGTGGCATGCTGGCGTATGCGTGGCTGCTGGAGTGCCCGGACCTGGTGGATCGTGCGGTTATCATTGGTGCGCCGGCGCGTCACAGCCCGTGGGCGATTGGTCTGAACACCGCGGCGCGTAACGCGATTCGTGCGGCGCCGGGTGGCGAGGGTCTGAAGGTTGCGCGTCAGATCGCGATGCTGAGCTATCGTAGCCCGGAGAGCTTCGCGCTGACCCAGAGCGGTTGGGGCACCCGTCGTCCGGGCACCCCGGACATTACCACCTACCTGGAGCACCAGGGTGAAAAACTGAGCACCCGTTTCTGCGAGCGTAGCTATCTGGCGCTGACCGGCGCGATGGACCGTTTTCAACCGACCGATGCGGAACTGCGTAGCATCCGTGTGCCGGTTCTGGTGGTTGGTATTAGCAGCGATGTGCTGTACCCGCCGGCGGAAGTGCGTACCTATGCGGGTCTGCTGCCGCGTGGCCAGTACCTGGAACTGCAAAGCCCGCACGGCCATGATGCGTTCCTGATCGATCCGCAGGGTCTGCCGGAAGCGGCGGCGGCGTTCCTGCACGGTGCGTAActcgagCAACCTGGAGGCGGGCGCAGGCCCGCCTTTTaagctt

> metYX_CM_OQ291224

gaattcTTTATTCTTGACACTAGTCGGCCAAAATGATATAATACCTGAGTTTAACTTTAAGAGAGGTATATATTACCATGGGTAGCAAGAACTACCGTTTCGAGACCCTGCAAGTGCACGGTGGCCAAGAAGTTGACCCGACCACCAACAGCCGTGCGGTGCCGATCTACCAGACCAGCAGCTATGTTTTTAACAGCGCGGAGCATGGTGCGAACCTGTTCGCGCTGAAGGAATTTGGCAACATCTATACCCGTATTATGAACCCGACCAGCGACGTTTTCGAGAAACGTATGGCGGCGCTGGAAGGTGGCGTGGCGGCGGTTGCGACCGCGAGCGGTCAGGCGGCGCAATTCCTGGCGCTGAACAACTTTCTGAGCGTGGGCGATAACTTCGTTACCAGCCCGTTTCTGTACGGTGGCAGCTATAACCAATTCAAAGTGAGCTTTAAACGTATCGGTATTGAGGCGCGTTTTGCGAAGAGCGACAAAGTTGACGATCTGGCGGCGGAAATCAACGATAAGACCAAAGCGATCTACGTGGAGACCATTGGCAACCCGGAGTTCAACGTTCCGGACTTTGAAGCGATCGCGGCGCTGGCGAAGAAACACGACATTCCGCTGGTTGTGGATAACACCTTCGGTGCGGGTGGCTATCTGTGCCAGCCGATCAAGCACGGCGCGAACATTGTGACCAGCAGCGCGACCAAATGGATCGGTGGTCACGGCACCAGCATTGGTGGCATCATTGTTGATGGTGGCAACTACAACTGGGGTAACGGCAAGTTCCCGCAATTTAGCGAGCCGAGCGAAGGTTATCACGGCCTGAACTTCTGGGAGACCTTTGGTGACAACAACCCGCTGGGTCTGCCGAACATTGCGTTCGCGATTCGTGCGCGTGTGGAAGGTCTGCGTGATTTTGGCCCGGCGATCAGCCCGTTCAACAGCTTTCTGCTGCTGCAAGGTCTGGAGACCCTGAGCCTGCGTGTGCAACGTACCGTTGACAACGCGCTGGAGCTGGCGAAATGGCTGGAAGCGCACCCGAAGGTGAAAAGCGTTAACTATCCGGGTCTGACCAACAGCCCGTACCACGCGACCGCGAAGAAATATCTGACCCACGGTTTCGGTGGCGTGCTGAGCTTTGAGATTGAAGGCGATAAGGAGACCGCGAGCAACTTTATCAACAACCTGGAACTGATTAGCCACCTGGCGAACGTTGGTGACGCGAAAACCCTGATCATTCAGCCGAGCGCGACCACCCACCAGCAACTGAGCGATGAAGCGCAAATTGCGGCGGGTGTGACCCCGAGCCTGCTGCGTATTAGCAGCGGCATCGAGCACATTGAAGACCTGAAGGCGGATCTGACCGCGGCGTTCGATAAAATCTAAGGATCCAGAGGTATATATTAATGAACCTGCAAAGCCCGCACCTGACCATCGAGATGACCCAGGAAATTTTCTACTGCCAAGAGGCGCTGAGCCTGGAGAGCGGCGAGAGCTTCCCGGAATTTCAACTGAGCTTTACCACCCAGGGCCAACTGAACGCGAACAAGGACAACGTGATCTGGGTTCTGCACGCGCTGACCGGTGATGCGAACCCGCACGAGTGGTGGAGCGGTCTGATCGGCGAAGACAAGTTCTTTGATCCGAGCAAATATTTCATTGTGTGCGCGAACTTTCTGGGTAGCTGCTACGGCAGCACCCAGCCGCTGAGCAACAACCCGAACAACGGTAAACCGTACTATTACGACTTCCCGAACATCACCACCCGTGACATTGCGAGCGCGCTGGATAAGCTGCGTATCCACCTGGGCCTGGAGAAAATCAACACCGTGATTGGTGGCAGCCTGGGTGGCCAAGTGGGTCTGGAATGGGCGGTTAGCCTGGGCGAGAAGCTGGAAAACGCGATCATTGTTGCGAGCAACGCGAAAGCGAGCCCGTGGATCATTGGTTTTAACGAGACCCAGCGTATGGCGATCGAAAGCGATAGCACCTGGGGCAAGACCCAACCGGAGGCGGGTAAGAAAGGCCTGGAAACCGCGCGTGCGATTGGTATGCTGAGCTATCGTCACCCGATGACCTTCCTGCAAAACCAAAGCGAGACCGAGGAAAAGCGTGACGATTTTAAAATCAGCAGCTATCTGCGTTACCAGGGCCTGAAGCTGGCGAACCGTTTCAACGCGATGAGCTACTGGATTCTGAGCAAAGCGATGGACAGCCACGATATTGGTCGTGGTCGTGGTGGCACCCCGGTGGCGCTGAGCAACATCAAGTGCAAAGTGCTGAGCATCGGTGTTGACACCGATATTCTGTTTACCAGCGAGGAAAGCCGTTATATTAGCAAGCACGTTCCGAAAGGCACCTATCGTGAGATCAGCAGCATTTACGGCCACGACGCGTTCCTGATCGAGTATGAACAGCTGCAATACATTCTGAAGAGCTTCTACCTGGAAAACAACGGCTAACTCGAGCAACCTGGAGGCGGGCGCAGGCCCGCCTTTTaagctt

> metYX_LI_OQ291225

gaattcTTTATTCTTGACACTAGTCGGCCAAAATGATATAATACCTGAGTTTAACTTTAAGAGAGGTATATATTACCATGGGTCCGCGTAACTATAAACCGGAGACCATTGCGCTGCACGGTGGCCAGAGCCCGGACCCGAGCACCCTGAGCCGTGCGGTGCCGATTTACCAAACCACCAGCTATGTTTTCAAGAACACCGAGCACGCGGCGAAACTGTTTGGTCTGCAAGAGTTCGGCAACATCTACACCCGTATTATGAACCCGACCACCGATGTTCTGGAGCAACGTATCGCGGCGCTGGAAGGTGGCGTGGCGGCGCTGGCGACCGCGAGCGGTCAGGCGGCGGAAACCCTGGCGCTGCTGAACATCGTGGAGGCGGGCCAAGAAATTGTTGCGAGCAGCAGCCTGTACGGTGGCACCTATAACCTGCTGCACTATACCTTTCCGAAGCTGGGTATCAAAGTGCACTTCGTTGACCCGAGCGATCCGGAGAACTTTCGTAAGGCGGTTAACGACAAAACCCGTGCGTTTTACGCGGAAACCCTGGGCAACCCGAAGCTGGATACCCTGAACCTGGAGGCGATTGCGAAAGTGGCGCACGACAGCGAAGTTCCGCTGATCATTGATAACACCCTGCCGAGCCCGTACCTGGTTAACCCGATCGAGCACGGTGCGGACATTGTGGTTCACAGCCTGACCAAGTTCCTGGGTGGTCACGGCACCAGCATCGGTGGCATCATTGTGGACAGCGGCAAATTTAACTGGGGTAACGGCAAGTTCAAAAACTTTACCGAACCGGACCCGAGCTATCACGGTCTGAAGTTCTGGGAAGTGTTCGGCAAATTTGAACCGTTCGGTGGCGTTAACATCGCGTACATCATTAAGGCGAAAGTGCAGGGTCTGCGTGATATGGGCGCGAGCATCAGCCCGTTTAACGCGTGGCAGATTCTGCAAGGTGTTGAGACCCTGCCGCTGCGTATGCGTAAACACAGCGAGAACGCGCTGGCGGTGGCGGAATATCTGACCAAGCACACCAAAGTGAGCTGGGTTAACTACCCGGGTCTGAAGATGGACAAAAACTACAGCCTGGCGAAGAAATATCACAAGAAAGATCTGTACGGCGCGATCCTGGGTTTCGGCATTAAGGGTGGCGCGGTGGAGGCGAAGAAATTTATCGACGGTCTGGAACTGTTCAGCCTGCTGGCGAACGTGGGCGATGCGAAAAGCCTGGTTATTCACCCGGCGAGCACCACCCACCAGCAACTGACCCCGGAGGAACAACTGAGCGCGGGTGTTACCCCGGACTTCGTGCGTCTGAGCGTTGGCCTGGAGAACATCGAAGACATTCTGTTTGATCTGGAGGAAGCGCTGAAGAAAGTGTAAGGATCCAGAGGTATATATTAATGAACGAGACCGGTAGCATCGGCATCATTGAAACCAAGTACGCGGAGTTCAAAGAACTGATCCTGAACAACGGTAGCGTGCTGAGCCCGGTGGTTATTGCGTACGAGACCTATGGCACCCTGAGCAGCAGCAAGAACAACGCGATCCTGATTTGCCATGCGCTGAGCGGTGATGCGCATGCGGCGGGTTATCACAGCGGCAGCGATAAGAAACCGGGTTGGTGGGACGATTACATTGGTCCGGGCAAGAGCTTCGACACCAACCAGTATTTTATCATTTGCAGCAACGTTATCGGTGGCTGCAAAGGTAGCAGCGGCCCGCTGAGCATTCATCCGGAGACCAGCACCCCGTATGGTAGCCGTTTCCCGTTTGTGAGCATCCAGGATATGGTTAAGGCGCAAAAGCTGCTGGTTGAGAGCCTGGGTATTGAAAAACTGTTCTGCGTTGCGGGTGGCAGCATGGGTGGCATGCAAGCGCTGGAATGGAGCATCGCGTATCCGAACAGCCTGAGCAACTGCATTGTTATGGCGAGCACCGCGGAGCACAGCGCGATGCAGATCGCGTTTAACGAAGTTGGTCGTCAAGCGATTCTGAGCGACCCGAACTGGAAGAACGGCCTGTACGATGAGAACAGCCCGCGTAAAGGTCTGGCGCTGGCGCGTATGGTGGGTCACATCACCTATCTGAGCGACGATAAGATGCGTGAAAAATTCGGTCGTAACCCGCCGCGTGGCAACATCCTGAGCACCGACTTTGCGGTTGGTAGCTACCTGATTTATCAGGGCGAGAGCTTCGTGGACCGTTTTGATGCGAACAGCTACATCTATGTTACCAAGGCGCTGGACCACTACAGCCTGGGCAAGGGCAAAGAACTGACCGCGGCGCTGAGCAACGCGACCTGCCGTTTCCTGGTGGTTAGCTACAGCAGCGATTGGCTGTATCCGCCGGCGCAAAGCCGTGAGATTGTGAAGAGCCTGGAAGCGGCGGACAAACGTGTGTTCTACGTTGAGCTGCAAAGCGGTGAAGGCCACGATAGCTTTCTGCTGAAGAACCCGAAACAAATCGAGATTCTGAAAGGTTTTCTGGAAAACCCGAACTAACTCGAGCAACCTGGAGGCGGGCGCAGGCCCGCCTTTTaagctt

>metAB

gaattcTTTATTCTTGACACTAGTCGGCCAAAATGATATAATACCTGAGTTTAACTTTAAGAGAGGTATATATTACCATGGGTACGCGTAAACAGGCCACCATCGCAGTGCGTAGCGGGTTAAATGACGACGAACAGTATGGTTGCGTTGTCCCACCGATCCATCTTTCCAGCACCTATAACTTTACCGGATTTAATGAACCGCGCGCGCATGATTACTCGCGTCGCGGCAACCCAACGCGCGATGTGGTTCAGCGTGCGCTGGCAGAACTGGAAGGTGGTGCCGGTGCAGTGCTCACCAATACCGGCATGTCAGCCATCCATCTGGTAACCACCGTCTTTTTGAAACCTGGCGATCTGCTGGTTGCGCCGCACGACTGCTACGGCGGTAGCTATCGCCTGTTCGACAGTCTGGCGAAACGCGGTTGCTATCGCGTGTTGTTTGTTGATCAAGGCGATGAACAGGCAATACGGGCAGCGCTGGCAGAAAAACCCAAACTGGTACTGGTAGAAAGCCCAAGTAATCCATTGTTACGCGTCGTGGATATTGCGAAAATCTGCCATCTGGCAAGGGAAGTCGGGGCGGTGAGCGTGGTGGATAACACCTTCTTAAGCCCGGCATTACAAAATCCGCTGGCATTAGGTGCCGATCTGGTGTTGCATTCATGCACGAAATATCTGAACGGTCACTCAGACGTAGTGGCCGGCGTGGTGATTGCTAAAGACCCGGACGTTGTCACTGAACTGGCCTGGTGGGCAAACAATATTGGCGTGACGGGCGGCGCGTTTGACAGCTATCTGCTGCTACGTGGGTTGCGAACGCTGGTGCCGCGTATGGAGCTGGCGCAGCGCAACGCGCAGGCGATTGTGAAATACCTGCAAACCCAGCCGTTGGTGAAAAAACTGTATCACCCGTCGTTGCCGGAAAATCAGGGGCATGAAATTGCCGCGCGCCAGCAAAAAGGCTTTGGCGCAATGTTGAGTTTTGAACTGGATGGCGATGAGCAGACGCTGCGTCGTTTCCTGGGCGGGCTGTCGTTGTTTACGCTGGCGGAATCATTAGGGGGAGTGGAAAGTTTAATCTCTCACGCCGCAACCATGACACATGCAGGCATGGCACCAGAAGCGCGTGCTGCCGCCGGGATCTCCGAGACGCTGCTGCGTATCTCCACCGGTATTGAAGATGGCGAAGATTTAATTGCCGACCTGGAAAATGGCTTCCGGGCTGCAAACAAGGGGTAAGGATCCAGAGGTATATATTAATGACGCGTAAACAGGCCACCATCGCAGTGCGTAGCGGGTTAAATGACGACGAACAGTATGGTTGCGTTGTCCCACCGATCCATCTTTCCAGCACCTATAACTTTACCGGATTTAATGAACCGCGCGCGCATGATTACTCGCGTCGCGGCAACCCAACGCGCGATGTGGTTCAGCGTGCGCTGGCAGAACTGGAAGGTGGTGCCGGTGCAGTGCTCACCAATACCGGCATGTCAGCCATCCATCTGGTAACCACCGTCTTTTTGAAACCTGGCGATCTGCTGGTTGCGCCGCACGACTGCTACGGCGGTAGCTATCGCCTGTTCGACAGTCTGGCGAAACGCGGTTGCTATCGCGTGTTGTTTGTTGATCAAGGCGATGAACAGGCAATACGGGCAGCGCTGGCAGAAAAACCCAAACTGGTACTGGTAGAAAGCCCAAGTAATCCATTGTTACGCGTCGTGGATATTGCGAAAATCTGCCATCTGGCAAGGGAAGTCGGGGCGGTGAGCGTGGTGGATAACACCTTCTTAAGCCCGGCATTACAAAATCCGCTGGCATTAGGTGCCGATCTGGTGTTGCATTCATGCACGAAATATCTGAACGGTCACTCAGACGTAGTGGCCGGCGTGGTGATTGCTAAAGACCCGGACGTTGTCACTGAACTGGCCTGGTGGGCAAACAATATTGGCGTGACGGGCGGCGCGTTTGACAGCTATCTGCTGCTACGTGGGTTGCGAACGCTGGTGCCGCGTATGGAGCTGGCGCAGCGCAACGCGCAGGCGATTGTGAAATACCTGCAAACCCAGCCGTTGGTGAAAAAACTGTATCACCCGTCGTTGCCGGAAAATCAGGGGCATGAAATTGCCGCGCGCCAGCAAAAAGGCTTTGGCGCAATGTTGAGTTTTGAACTGGATGGCGATGAGCAGACGCTGCGTCGTTTCCTGGGCGGGCTGTCGTTGTTTACGCTGGCGGAATCATTAGGGGGAGTGGAAAGTTTAATCTCTCACGCCGCAACCATGACACATGCAGGCATGGCACCAGAAGCGCGTGCTGCCGCCGGGATCTCCGAGACGCTGCTGCGTATCTCCACCGGTATTGAAGATGGCGAAGATTTAATTGCCGACCTGGAAAATGGCTTCCGGGCTGCAAACAAGGGGTAACTCGAGCAACCTGGAGGCGGGCGCAGGCCCGCCTTTTaagctt
